# Resting State Neural Networks at Complex Visual Hallucinations in Charles Bonnet Syndrome

**DOI:** 10.1101/2022.07.15.500190

**Authors:** Taha Hanoglu, Halil Aziz Velioglu, Behram Ali Salar, Sultan Yıldız, Zübeyir Bayraktaroglu, Burak Yulug, Lutfu Hanoglu

## Abstract

**Background:** Charles Bonnet syndrome (CBS) is a prototype phenomenon for investigating complex visual hallucination. Our research focuses on resting state neural networks features of CBS patients with a comparison of patients with equally matched visual loss and healthy subjects in order to investigate the mechanism behind complex visual hallucinations.

**Material and Methods:** Four CBS patients CBS(+), three patients with visual loss but no visual hallucinations CBS(-) and 15 healthy individuals (HS) undergo resting state fMRI recordings and their resting state data is analyzed for Default Mode Network (DMN) changes through dual regression analysis. Cognitive functions of the participants were also evaluated through Mini Mental State Examination and University of Miami - Parkinson’s Disease Hallucination Questionnaire (um-PDHQ)

**Results:** Although we found no difference in Default Mode Networks between CBS(-) and CBS(+), and between the CBS(-) and HC groups, we detected decreased connectivity in CBS(+) compared to the HC group especially in visual heteromodal association centers (bilateral lateral occipital gyrus, bilateral lingual gyrus, occipital pole, right medial temporal cortex, right temporo-occipital cortex) when left angular gyrus was selected as ROI.

Similarly, we detected decreased connectivity in CBS(+) compared to HC in right medial frontal gyrus, right posterior cingulate gyrus, left inferior temporal gyrus, right supramarginal gyrus, and right angular gyrus when selected right superior frontal gyrus as ROI. In contrast, increased connectivity was detected in CBS +compared to HC, in bilateral occipital poles, bilateral occipital fusiform gyrus, bilateral intracalcarine cortex, right lingual gyrus and precuneus regions when left medial temporal gyrus was selected as ROI.

**Conclusion:** Our findings suggest a combined mechanism in CBS related to increased internal created images caused by decreased visual external input causing visual hallucinations as well as impaired frontotemporal resource tracking system that together impair cognitive processing.

## INTRODUCTION

Charles Bonnet syndrome (CBS) is classically defined as the experience of complex visual hallucinations secondary to pure visual deficiency in the absence of any cognitive impairment. The majority of hallucinations generated during CBS are complex visual hallucinations (1).

The most widely accepted hypothesis concerning the mechanism underlying these hallucinations in CBS involves spontaneous indigenous activity due to input deficiency caused by bottom-up deafferentation. This deafferentation causes depression in primary visual pathways, although it also causes a hyperexcitability state, especially in visual association areas emerging with the so-called “cortical release phenomenon”, a similar entity to the phantom limb phenomenon (2).

The first part of the deafferentation hypothesis, depression in the primary visual areas, has been shown through almost all metabolic studies in the field (3–6). However, despite these valuable metabolic data, this concept is not by itself sufficient to explain the hallucination pathophysiology. This may be because low metabolism can result in a direct deafferentation effect. Moreover, this hypothesis fails to adequately answer the critical question of why most patients with comparable visual loss develop no hallucinations (7). The second part of the deafferentation hypothesis suggests hyperexcitability in visual association areas, although, in contrast to prior consistent data, the evidence from metabolic studies is highly variable (3–5).

There are also mixed results, including our own findings, indicating that both primary and secondary areas are involved simultaneously in the pathophysiology of visual hallucinations. For example, Ffytche suggested dysregulation of the tonic-burst activation switch of the thalamo-cortical circuit (8), while our own recent data suggest a metabolic change in a broader spectrum, containing the posterior cingulate and inferior frontal cortices (9). These findings together suggested Collerton et al. ‘s “perception and attention deficit model”, implying a top-down dysregulation of the attention network that offers a combined explanation of bottom-up perspective defect (deafferentation) with top-down attention channeling disorder (attention network disorder). Collerton et al. created an overall valid visual hallucination model, rather than one specific for CBS hallucinations, to identify the point at which the source control deficit occurs in order to understand the underlying mechanisms of hallucination (10).

The default mode network (DMN) is defined as a functional network activated during the resting state and inhibited while executing a task. Imaging studies have shown that the DMN consists of cortical structures closely related to limbic areas, such as the medial prefrontal cortex, posterior cingulate, and inferior lateral parietal cortex (11,12). Several studies have suggested that the DMN functions actively during image creation and modifies or transmits self-referenced information to a conscious level (13). Several researchers have also investigated the role of DMN in the generation of visual hallucinations in Parkinson’s Disease and Lewy Body Dementias (14–20).

In the light of these studies, we aimed to examine the resting state functional network changes in individuals with CBS, in counterparts with comparable visual disturbance who developed no visual hallucinations, and in healthy controls in order to elucidate the underlying mechanisms of such hallucinations in healthy individuals and CBS. We also investigated whether a disease-specific approach or a universal integrative approach is more appropriate for understanding the nature of hallucinations in CBS and other disease conditions. Instead of focusing on one region, such as the primary visual cortex or visual association areas, which have been shown not to be appropriate for a whole frame inducing visual hallucination, we aimed to establish the integrated view of visual hallucination pathophysiology described by Collerton et al. and analyzed the resting state features of the participants in order to acquire further supporting evidence or improvement for this model.

## MATERIAL AND METHOD

### Study Groups

CBS(+) group consists of 4 individuals with complex visual hallucinations after acquired visual loss. Whereas, CBS(-) group consists of 3 individuals with visual loss statistically peered to those of CBS(+) group but no visual hallucinations. Lastly, the healthy controls group consists of 15 statistically peered individuals without any visual loss or visual hallucinations.

### Participant Selection Criteria

While creating the CBS(+) and CBS(-) groups, 208 patients with a visual loss of 0.3 or below from Istanbul Medipol University Ophthalmology Clinic are contacted and questioned via telephone about visual hallucinations (21). Twelve of those patients had experienced visual hallucinations and 4 of those 12, who had been diagnosed as CBS, had given permission to a rs-fMRI recording. Similarly, 4 patients within the same margin of visual loss had given permission to a rs-fMRI recording. Thus, the CBS(+) and CBS(-) had been chosen for the study.

CBS(+), CBS(-) and healthy control groups are subjected to Mini-mental State Examination (MMSE) (22) in order to evaluate the general cognitive function of the participants. Besides, CBS(+) groups visual hallucination content, frequency, duration and its effect on the subject are also investigated through University of Miami Parkinson’s Disease Hallucination Questionnaire (um-PDHQ) (23). Except for these, the participants had been evaluated for mood disorders via Geriatric Depression Scale (GDS) (24).

### Materials that are used in the Experiment

The structural and functional magnetic resonance imaging (fMRI) data of the participants are recorded on 3-Tesla (T) Philips Achieva (Philips Medical Systems, Best, the Netherlands)

The raw data had been transformed from DICOM (Digital Imaging and Communications in Medicine) to the for the analysis necessary format of NIFTI (Neuroimaging Informatics Technology Initiative) via dcm2niix (ver. v1.0.20170724) program.

The preprocessing, independent component analysis (ICA), artifact rejection and dual regression analysis of resting state MRI data had been executed via FSL (FMRIB Software Library, Oxford, England; http://www.fmrib.ox.ac.uk.fsl)

### Functional Magnetic Resonance Imaging (fMRI) Analysis

#### Gathering the Data

All of the fMRI recordings of participants carried out standardly at resting state, as the person taking his place in the device and after routine calibrations and positioning controls had been done. The participants ought to stay still and eyes open as the recordings were taking place.

The recording parameters of functional data was as following: FOV (Field of View): 128 x 128 x 36 mm (RL x AP x FH), Volume number: 180, Voxel Size: 1.86 x 1.86 x 4 mm, TR (Repetition Time): 2000 ms. Whereas the recording parameters of T1 structural data was: FOV: 190 x 256 x 256 mm., Voxel size: 1×1×1 mm, TR: 8 ms.

#### Preprocessing

The analysis was carried out via FMRIB FSL (ver.6.0, https://www.fmrib.ox.ac.uk/fsl) software using Linux Mint 18.3 Sylvia operating system. The RAW recording data format of DICOM was transformed into the FSL software compatible Nifti format via dcm2niix (ver. v1.0.20170724) software tool.

The analysis begins with extracting brain tissue from other tissues at the anatomic data with the combination of BET (Brain Extraction Tool) command of FSL and fsl_anat script to optimize the application of bias-field correction. Secondly, all the functional resting state data equalized to the 150th volume of the recording (the middle point of the recording) and so, motion artifacts are corrected in 3 axes and 3 planes with the FSL FEAT command. The artifacts caused by head-neck motion, respiration and kardio-vascular system are eliminated using a high-pass filter of 0.01-0.1 Hz (the interval that the resting state functional networks operate). The Functional data is softened with a softening value of 3 in order to fit the data to normal distribution. Following, the functional data is aligned to their own structural data with FSL FLIRT command using a boundary-based registration (BBR) approach. Immediately after, the self-registered data is registered to MNI152 standard brain image with FSL FNIRT command with a nonlinear registration method. At this point, all of the functional data is registered to the standard anatomical plane using the transformation matrices created by the beforehand analysis in order to prepare the data for the group analysis. Lastly, the data underwent the independent component analysis using the FSL MELODC ICA commend.

#### Artifact Rejection

The components created by the ICA process are visualized and evaluated with their frequency and localization features with the FSLeyes tool. The components with a frequency of 0.1 Hz or higher, or that have a spike shaped or quickly changing frequency patterns are flagged during the evaluation. The components with common artifact patterns and with activity signals out of the gray matter are also flagged. Only the components with pure resting state activity are labeled as signals. The flagged independent components are rejected from the data with the fsl_regfilt command. After that the artifact free data is transferred to the standard plane with applywarp command.

### Dual Regression and Region of Interest (ROI) Analysis

In order to compare the artifact free functional data in a group level a two-level dual regression analysis method was used. The average components of each group and the subtractions of study groups one another for all dual combinations are calculated with FSL dual_regression command at 5000 permutations setting. The experiment matrix is prepared with FSL’s GLM tool as shown in Fig 1.

**Fig 1.**
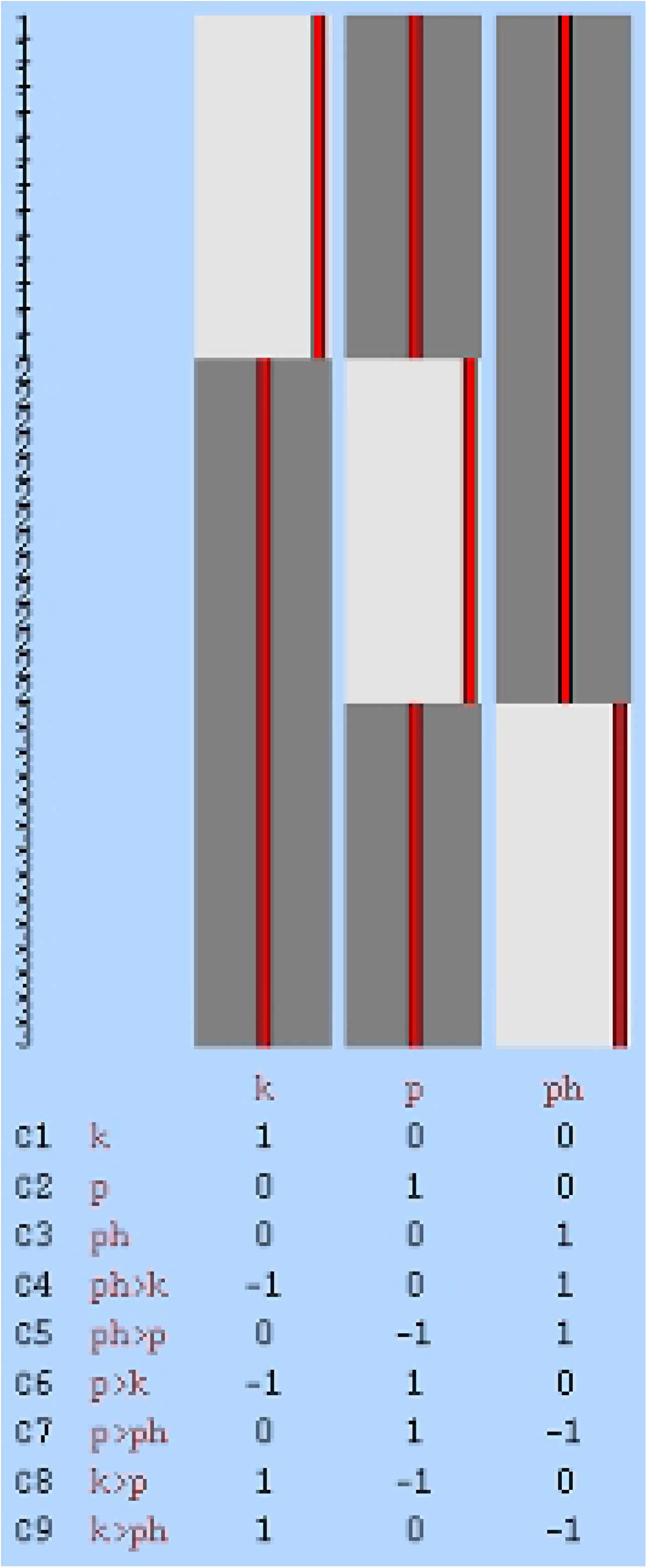
The experiment matrix that was prepared in FSL GLM tool is shown. C1- Health control group average activity C2- CBS(-) group average activity C3- CBS(+) group average activity C4- The subtraction of healthy control group from CBS(+) group C5- The subtraction of CBS(-) group from CBS(+) group C6- The subtraction of healthy control group from CBS(-) group C7- The subtraction of CBS(+) group from CBS(-) group C8- The subtraction of CBS(-) group from healthy control group C9- The subtraction of CBS(+) group from healthy control group

The components are compared one another using t-tests. The age differences are added to GLM matrices as co-parameter in order to reduce the effect of it. The significance level for the comparisons was regarded as p<0.05.

The dual regression analysis’ calculations of ICA components’ independent time series and spatial maps were realized with the “dual_regression” command in FSL. The ROIs used in dual regression analysis were chosen as the template that is suggested by Brown et al. for DMN [25]. This template consists of 10 different DMN components as following: paracingulate gyrus, precuneus, hippocampus, parahippocampal gyrus, medial temporal lobes, medial frontal gyri and, angular gyrus (Figs 2 and 3a,b,c). Thus, we were able to attain independent time series for every binary combination of CBS(+), CBS(-) and HS groups.

**Fig 2.**
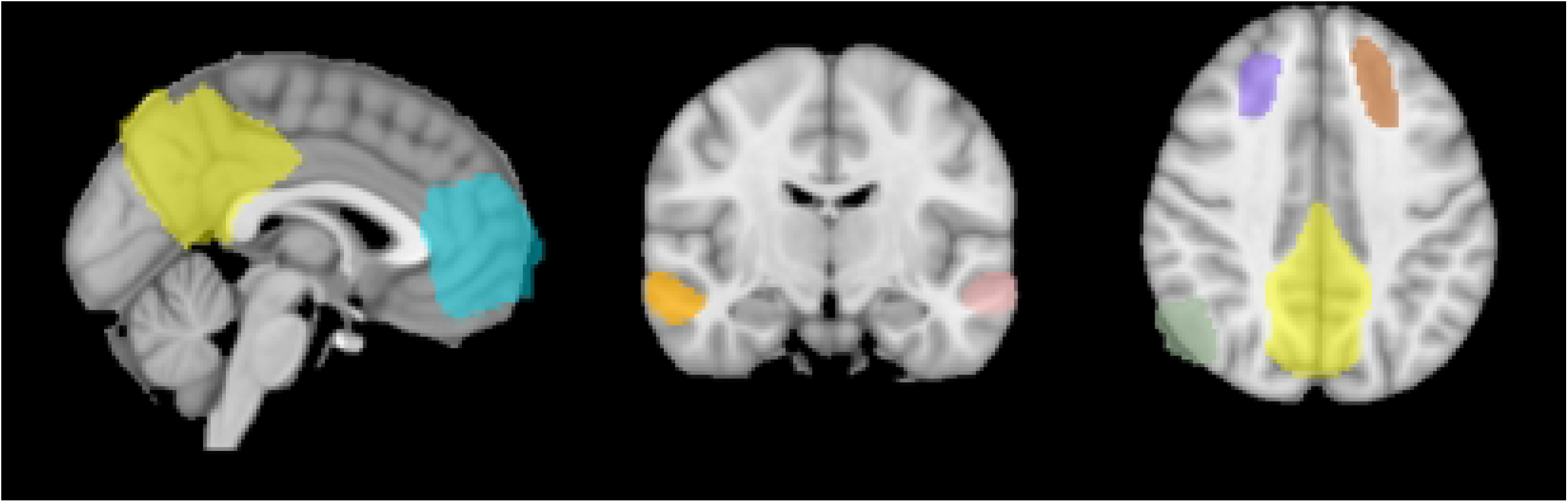
Brown et al. (2019) (25)DMN template. Chyan for paracingulate gyrus, yellow for precuneus, parliament blue for hippocampus, green for parahippocampal gyrus, pink for left medial temporal lobe, orange for right medial temporal lobe, purple for medial frontal gyrus, brown for right medial frontal gyrus and jade for angular gyrus

**Fig 3a,b,c.**
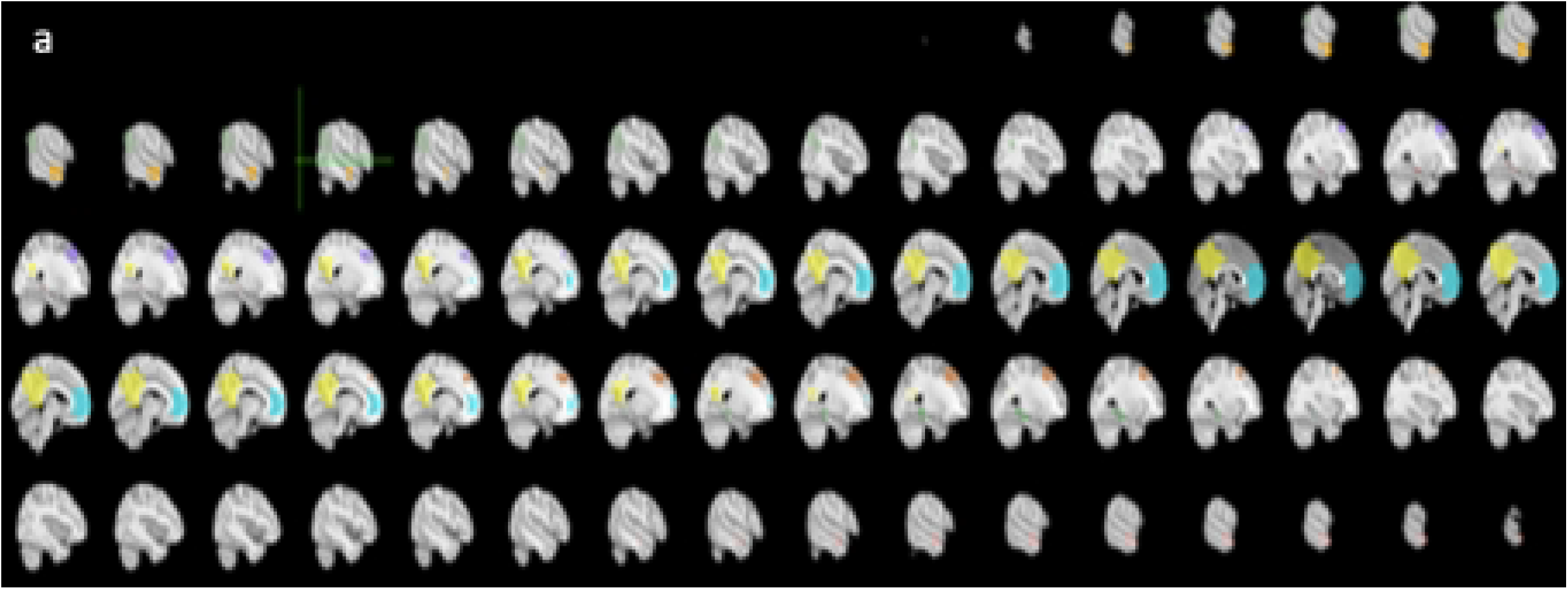
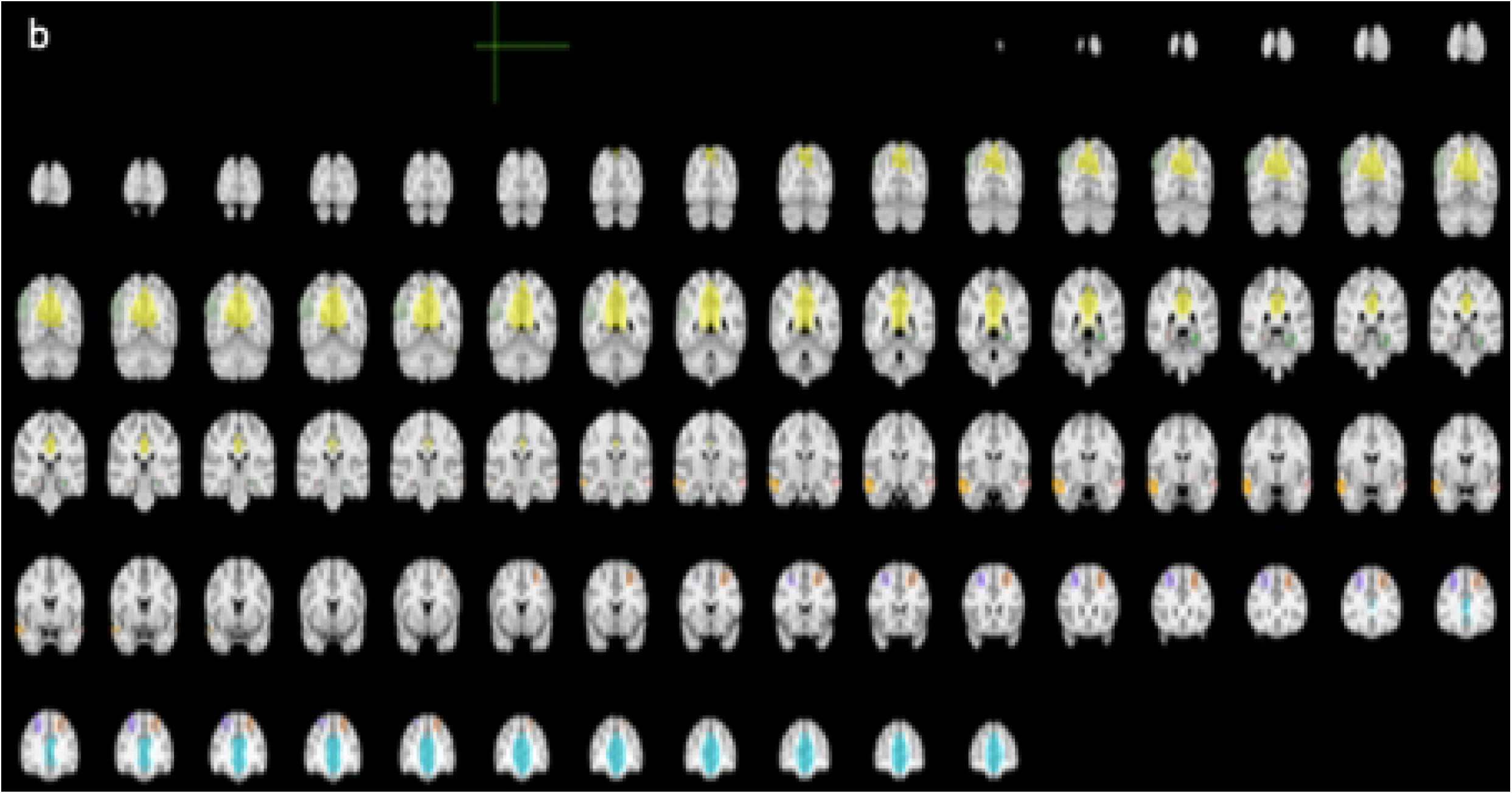
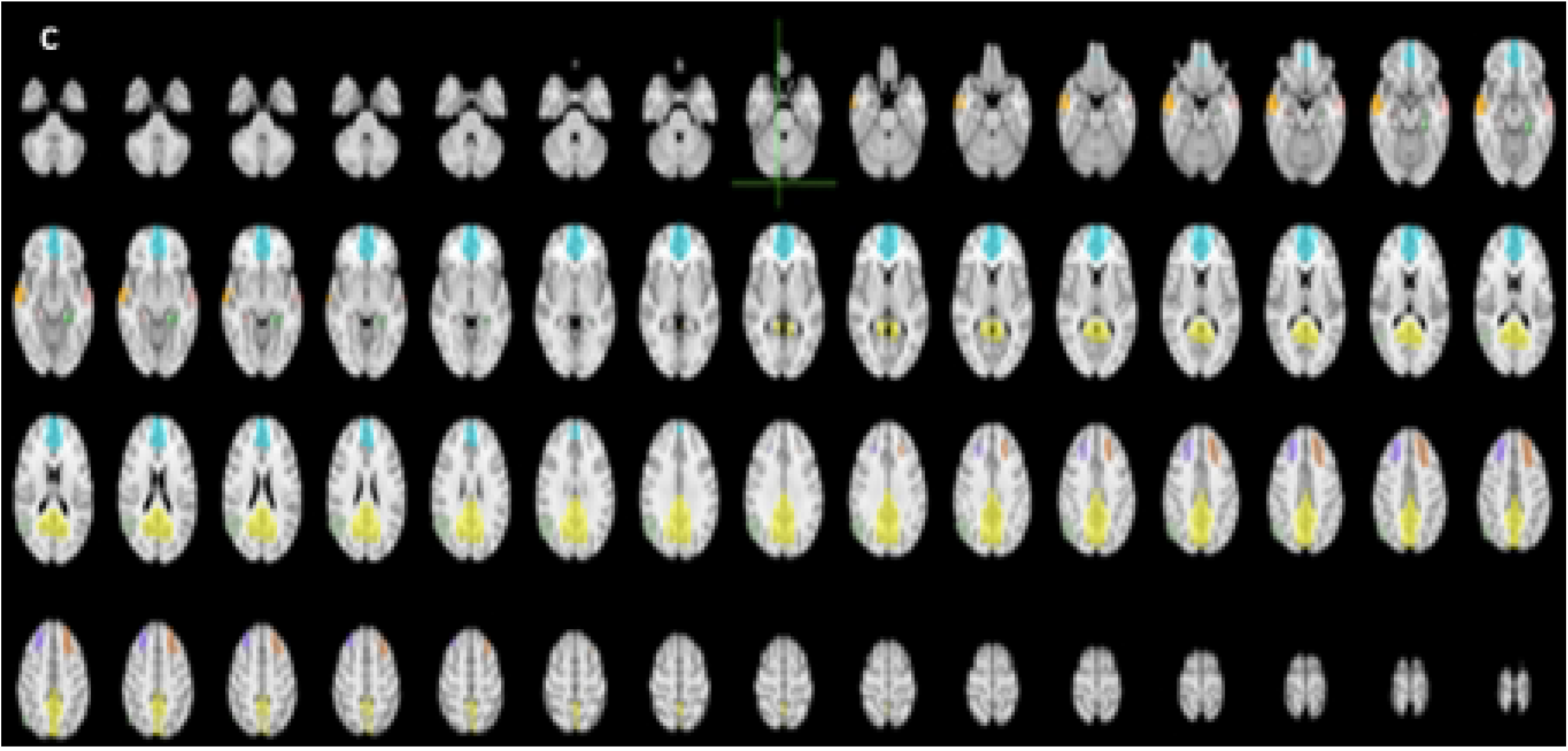
Brown et al. (2019) (25) DMN template. Chyan for paracingulate gyrus, yellow for precuneus, parliament blue for hippocampus, green for parahippocampal gyrus, pink for left medial temporal lobe, orange for right medial temporal lobe, purple for medial frontal gyrus, brown for right medial frontal gyrus and jade for angular gyrus

### Statistical Analysis

Groups (CBS(+), CBS(-) and HS) had been chosen as between-subject factor and age, sex, years of education, MMSE, GDS, um-PDHQ as in-between subject factors for the power spectrum analysis (quad vide Table 1).

**Table 1.**
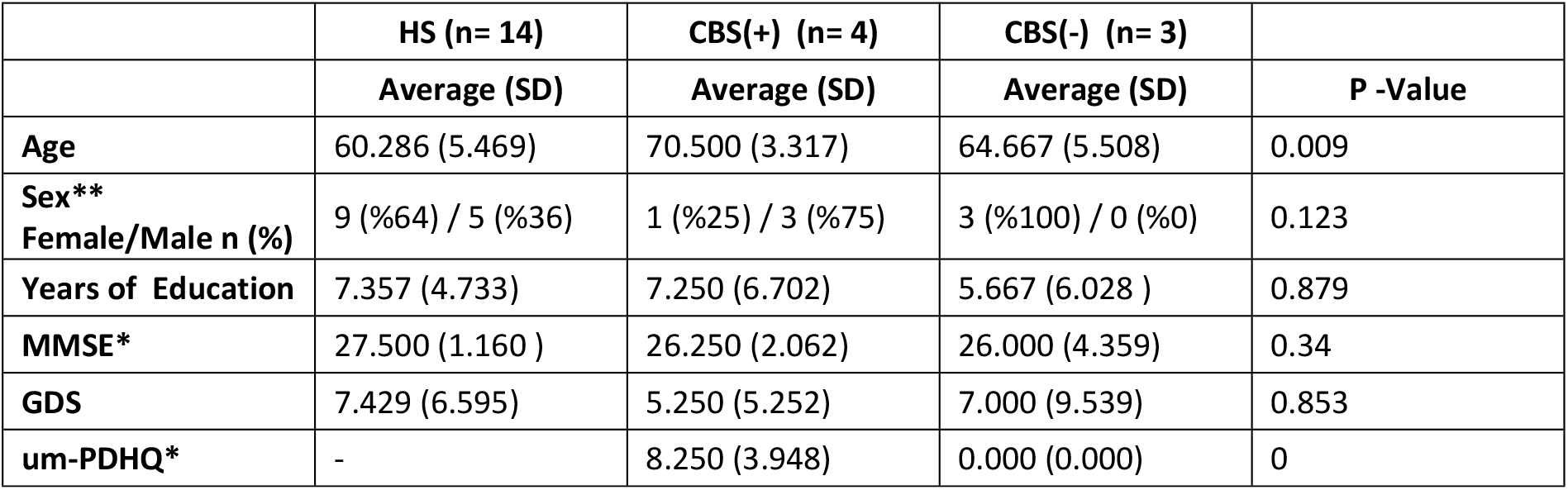
Table that demonstrates the demographics of study groups. One way Anova test was applied for the normally distributed parametric variables, whereas Kruskal-wallis test was applied for the parametric variables that distributed not normally (shown with “*”) and Chi-square test was applied for the non-parametric variables (shown with “*”).

The JASP software 0.14.1.0 version had been used for the statistical analysis (26). P < 0.05 accepted as the statistical significance level. Anova test was used to compare the parametric variables that showed a normal distribution. Kruskal-Wallis test was used to compare the parametric variables that did not show a normal distribution (shown with “*” in table 1). Chi-square test was used to compare the non-parametric variables (shown with “**” in table 1).

Kurtosis and skewness values were imposed on detecting the distribution features of the variables. The kurtosis and skewness values between (+ 2) and (−2) were accepted as normally distributed as George et al. suggested (27).

The Hochberg GT2 test was applied for the values that were statistically significant according to Anova and Kruskal-Wallis tests as a post-hoc test because of the immense subject difference in between the study groups.

### Ethical Committee Approval

The Ethical Committee of Istanbul Medipol University accepted our research with the number of Ethical Report: E-10840098-772.02-2331 at 25.05.2021.

## RESULTS

### The Demographics of Study Groups

Sex, years of education, MMSE, GDS variables showed no statistical significance, whereas age, um-PDHQ variables showed statistical significance as the demographics of the participants were investigated. Post-hoc analysis was applied to investigate the reason behind these statistical significances. The reason for statistical significance of the age variable was determined as HS group. On the other hand, all of the groups were determined to be statistically significant for the um-PDHQ variable as expected.

### Results of fMRI Analysis

There was no statistical significance to report of the fMRI analysis for DMN areas as regions of interest in the resting state data between the CBS(-) and CBS(+), and also between CBS(-) and HS. Whereas, there are several statistical significance for the same fMRI analysis between CBS(+) and HS. Firstly, we had detected a decreased functional connectivity between following brain regions; bilateral lateral occipital gyri, bilateral lingual gyri, occipital pole, right medial temporal cortex, right temporo-occipital cortex and the ROI in the left angular gyrus (see below Fig 4a,b,c and for the cluster values see table 2) in CBS(+) group in comparison to HS group.

**Fig 4a,b,c.**
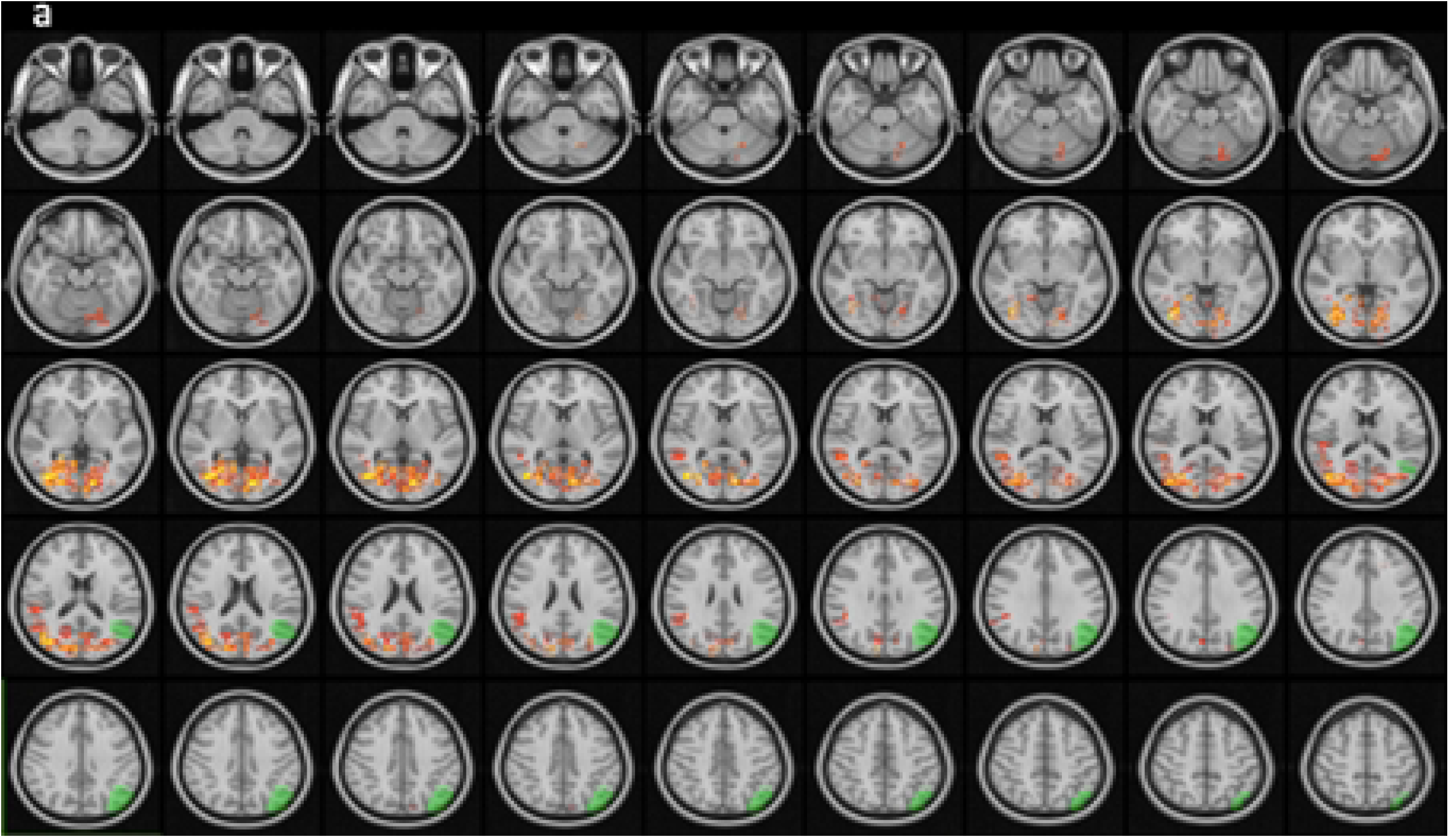
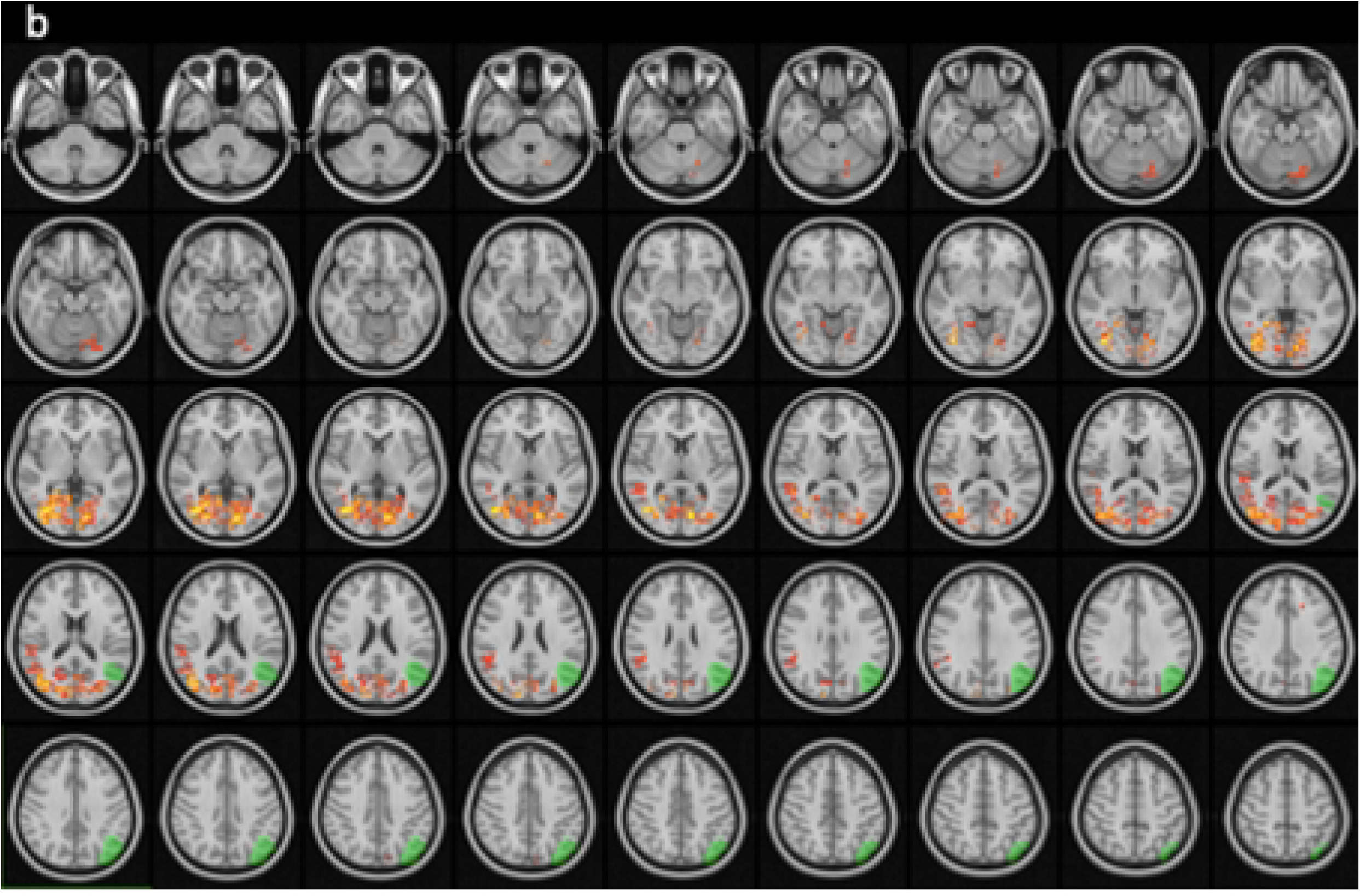
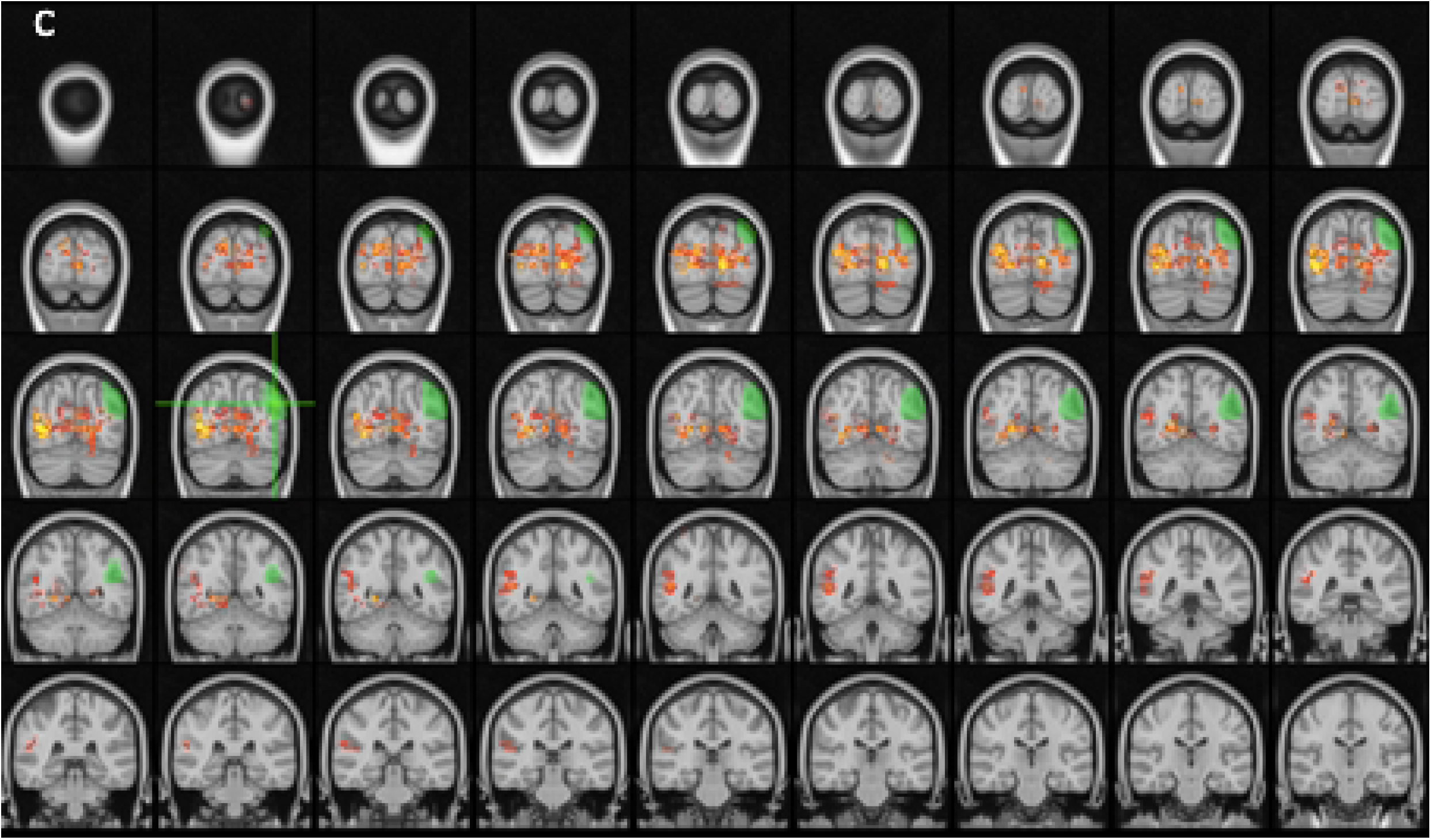
The areas with statistical-significantly decreased functional connectivity between ROI in left angular gyrus and bilateral lateral occipital gyri, bilateral lingual gyri, occipital pole, right medial temporal cortex and right temporo-occipital cortex, when the time series of HS group and CBS(+) group are compared with one way t-test in dual regression analysis (p<0,05).

**Table 2.**
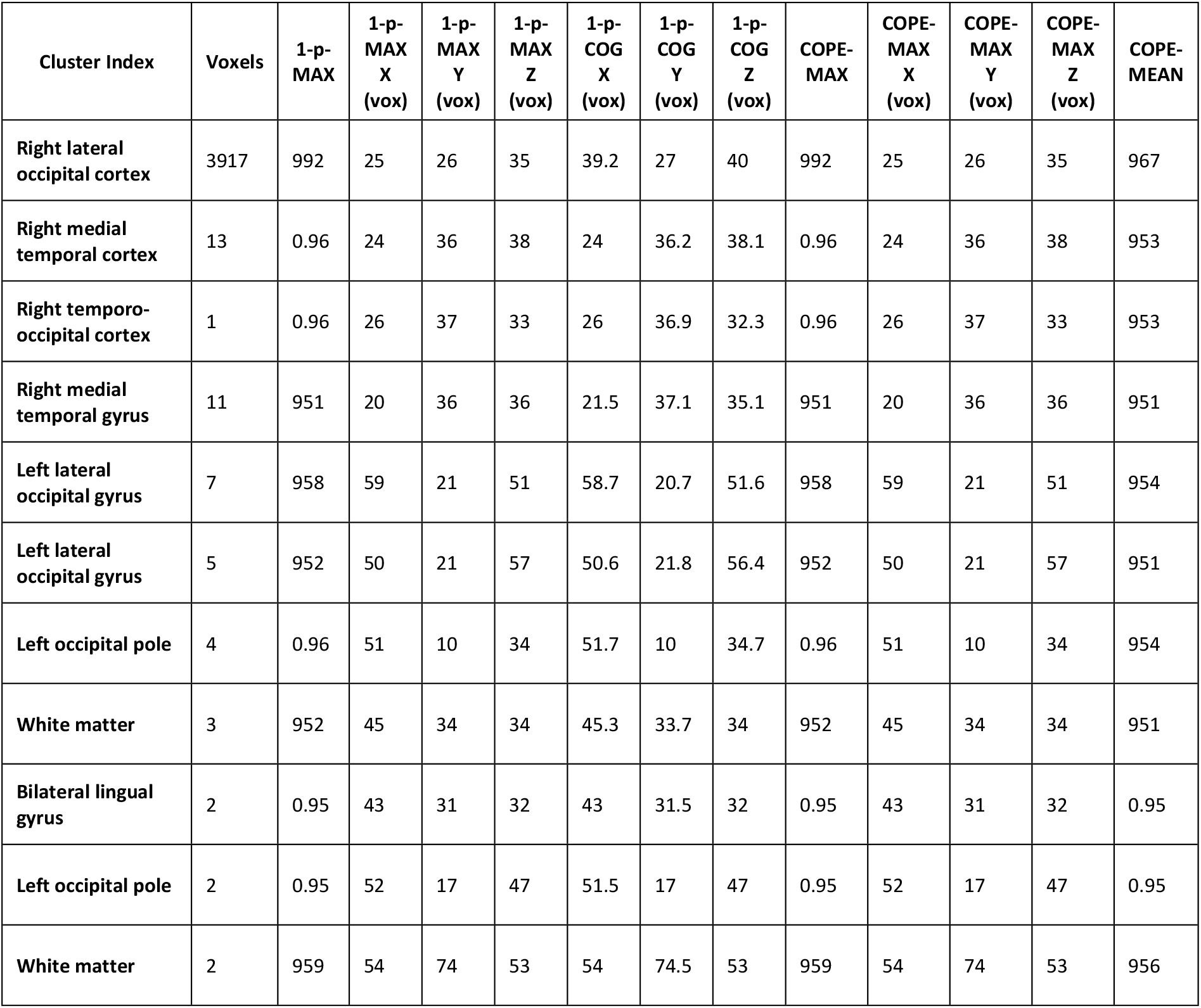
The cluster coordinates with statistical-significantly decreased functional connectivity between ROI in left angular gyrus and bilateral lateral occipital gyri, bilateral lingual gyri, occipital pole, right medial temporal cortex and right temporo-occipital cortex, when the time series of HS group and CBS(+) group are compared with one way t-test in dual regression analysis (p<0,05)

| Cluster Index | Voxels | 1-p-MAX | 1-p-MAX X (vox) | 1-p-MAX Y (vox) | 1-p-MAX Z (vox) | 1-p-COG X (vox) | 1-p-COG Y (vox) | 1-p-COG Z (vox) | COPE-MAX | COPE-MAX X (vox) | COPE-MAX Y (vox) | COPE-MAX Z (vox) | COPE-MEAN |
| --- | --- | --- | --- | --- | --- | --- | --- | --- | --- | --- | --- | --- | --- |
| Right lateral occipital cortex | 3917 | 992 | 25 | 26 | 35 | 39.2 | 27 | 40 | 992 | 25 | 26 | 35 | 967 |
| Right medial temporal cortex | 13 | 0.96 | 24 | 36 | 38 | 24 | 36.2 | 38.1 | 0.96 | 24 | 36 | 38 | 953 |
| Right temporo-occipital cortex | 1 | 0.96 | 26 | 37 | 33 | 26 | 36.9 | 32.3 | 0.96 | 26 | 37 | 33 | 953 |
| Right medial temporal gyrus | 11 | 951 | 20 | 36 | 36 | 21.5 | 37.1 | 35.1 | 951 | 20 | 36 | 36 | 951 |
| Left lateral occipital gyrus | 7 | 958 | 59 | 21 | 51 | 58.7 | 20.7 | 51.6 | 958 | 59 | 21 | 51 | 954 |
| Left lateral occipital gyrus | 5 | 952 | 50 | 21 | 57 | 50.6 | 21.8 | 56.4 | 952 | 50 | 21 | 57 | 951 |
| Left occipital pole | 4 | 0.96 | 51 | 10 | 34 | 51.7 | 10 | 34.7 | 0.96 | 51 | 10 | 34 | 954 |
| White matter | 3 | 952 | 45 | 34 | 34 | 45.3 | 33.7 | 34 | 952 | 45 | 34 | 34 | 951 |
| Bilateral lingual gyrus | 2 | 0.95 | 43 | 31 | 32 | 43 | 31.5 | 32 | 0.95 | 43 | 31 | 32 | 0.95 |
| Left occipital pole | 2 | 0.95 | 52 | 17 | 47 | 51.5 | 17 | 47 | 0.95 | 52 | 17 | 47 | 0.95 |
| White matter | 2 | 959 | 54 | 74 | 53 | 54 | 74.5 | 53 | 959 | 54 | 74 | 53 | 956 |

Secondly, we had detected a decreased functional connectivity between following brain regions; right medial frontal gyrus, right posterior cingulate gyrus, right parietal lobe, right inferior temporal gyrus, right supramarginal gyrus and the ROI in the right superior frontal gyrus (see below Fig 5a,b,c and for the cluster values see table 3) in CBS(+) group in comparison to HS group.

**Fig 5a,b,c.**
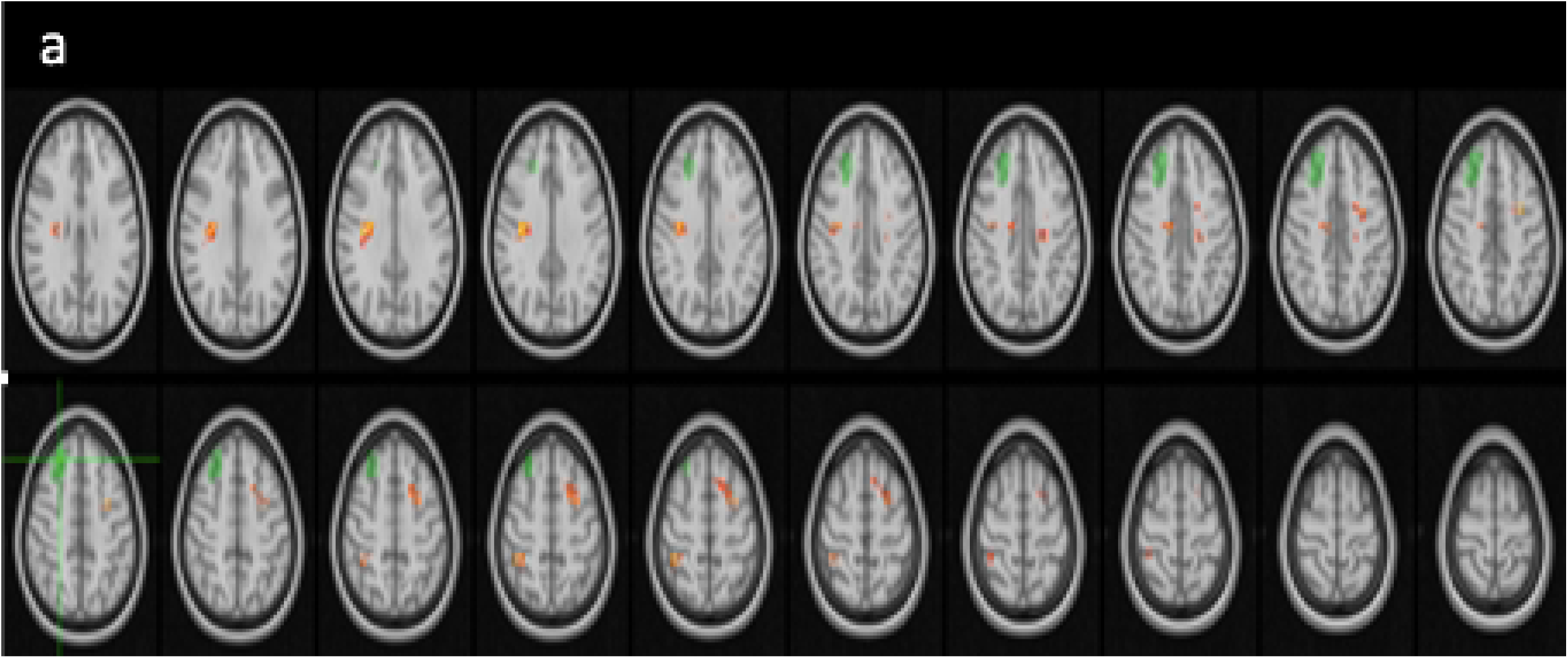
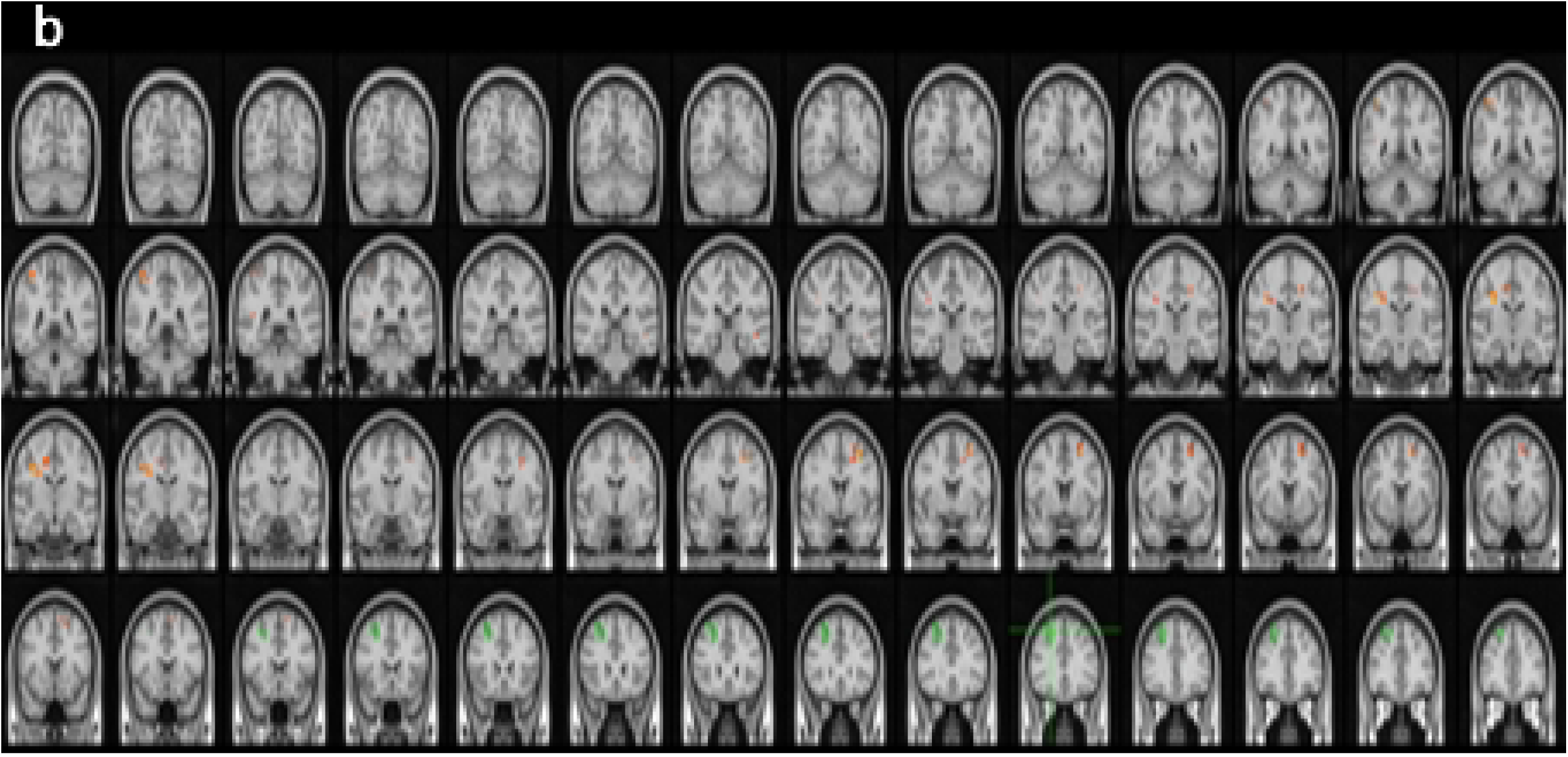
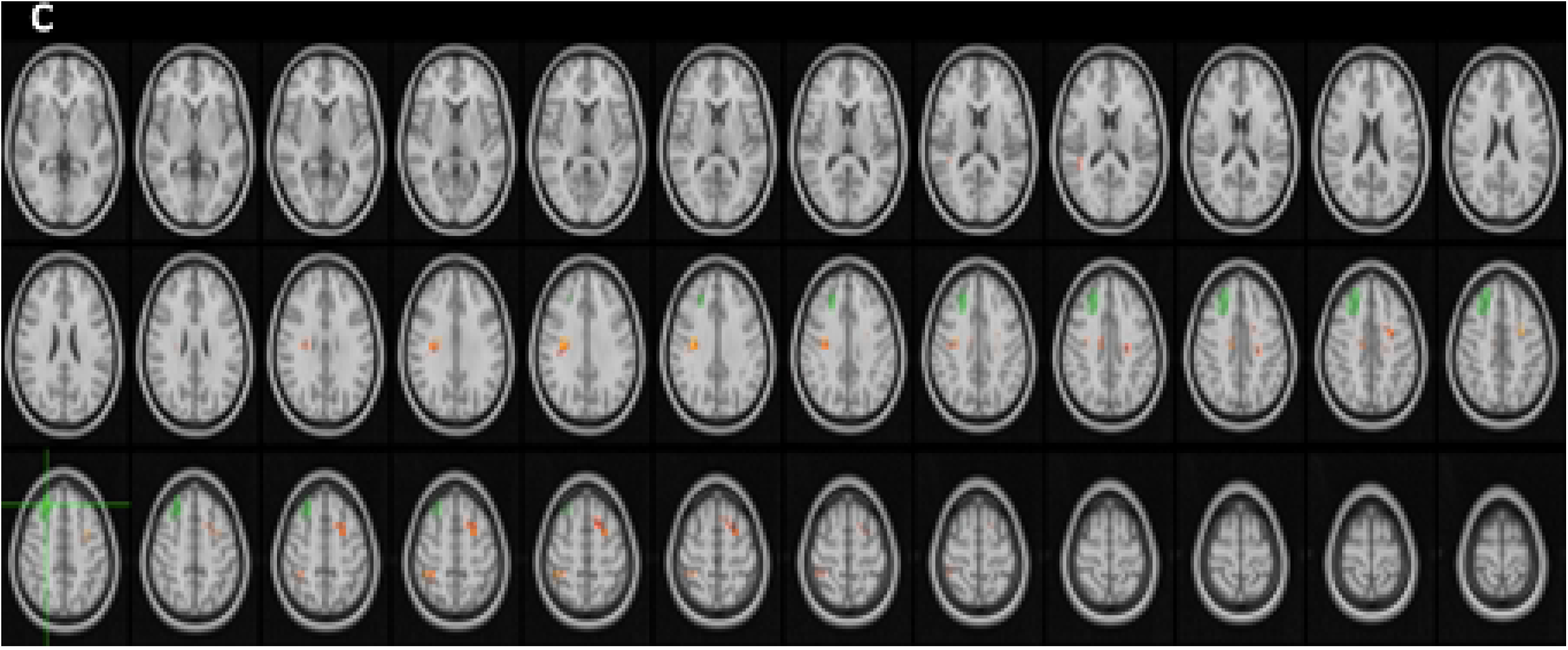
The areas with statistical-significantly decreased functional connectivity between ROI in right superior frontal gyrus and right medial frontal gyrus, right posterior cingulate gyrus, right superior parietal lobe, left inferior temporal gyrus, right supramarginal gyrus, right angular gyrus, when the time series of HS group and CBS(+) group are compared with one way t-test in dual regression analysis (p<0,05).

**Table 3.** The cluster coordinates with statistical-significantly decreased functional connectivity between ROI in right superior frontal gyrus and right medial frontal gyrus, right posterior cingulate gyrus, right superior parietal lobe, left inferior temporal gyrus, right supramarginal gyrus, right angular gyrus, when the time series of HS group and CBS(+) group are compared with one way t-test in dual regression analysis (p<0,05).

| Cluster Index | Voxels | 1-p-MAX | 1-p-MAX X (vox) | 1-p-MAX Y (vox) | 1-p-MAX Z (vox) | 1-p-COG X (vox) | 1-p-COG Y (vox) | 1-p-COG Z (vox) | COPE-MAX | COPE-MAX X (vox) | COPE-MAX Y (vox) | COPE-MAX Z (vox) | COPE-MEAN |
| --- | --- | --- | --- | --- | --- | --- | --- | --- | --- | --- | --- | --- | --- |
| Right medial frontal gyrus | 155 | 972 | 59 | 63 | 59 | 55.9 | 65.4 | 62.1 | 972 | 59 | 63 | 59 | 958 |
| Right posterior cingulate gyrus | 153 | 975 | 27 | 56 | 53 | 28.4 | 54.9 | 52.5 | 975 | 27 | 56 | 53 | 0.96 |
| Right superior parietal lobe | 56 | 0.97 | 25 | 40 | 63 | 25.8 | 42 | 64 | 0.97 | 25 | 40 | 63 | 959 |
| Right posterior cingulate gyrus | 39 | 964 | 38 | 55 | 57 | 37.9 | 55.9 | 57 | 964 | 38 | 55 | 57 | 957 |
| White matter | 23 | 0.96 | 56 | 52 | 56 | 56 | 52.6 | 56.5 | 0.96 | 56 | 52 | 56 | 955 |
| Left inferior temporal gyrus | 8 | 961 | 67 | 47 | 32 | 67.6 | 48 | 32.4 | 961 | 67 | 47 | 32 | 955 |
| Right supramarginal gyrus | 6 | 958 | 24 | 44 | 44 | 24 | 44.3 | 43.7 | 958 | 24 | 44 | 44 | 953 |
| Right angular gyrus | 4 | 954 | 22 | 40 | 44 | 22.7 | 40 | 44.2 | 954 | 22 | 40 | 44 | 952 |

Lastly, we had detected an increased functional connectivity between following brain regions; bilateral occipital poles, bilateral occipital fusiform gyri, bilateral intra-calcarine cortices, right lingual gyrus, precuneus, medial temporal gyrus and the ROI in the left medial temporal gyrus and (see below Fig 6a,b,c and for the cluster values see table 4) in CBS(+) group in comparison to HS group.

**Fig 6a,b,c.**
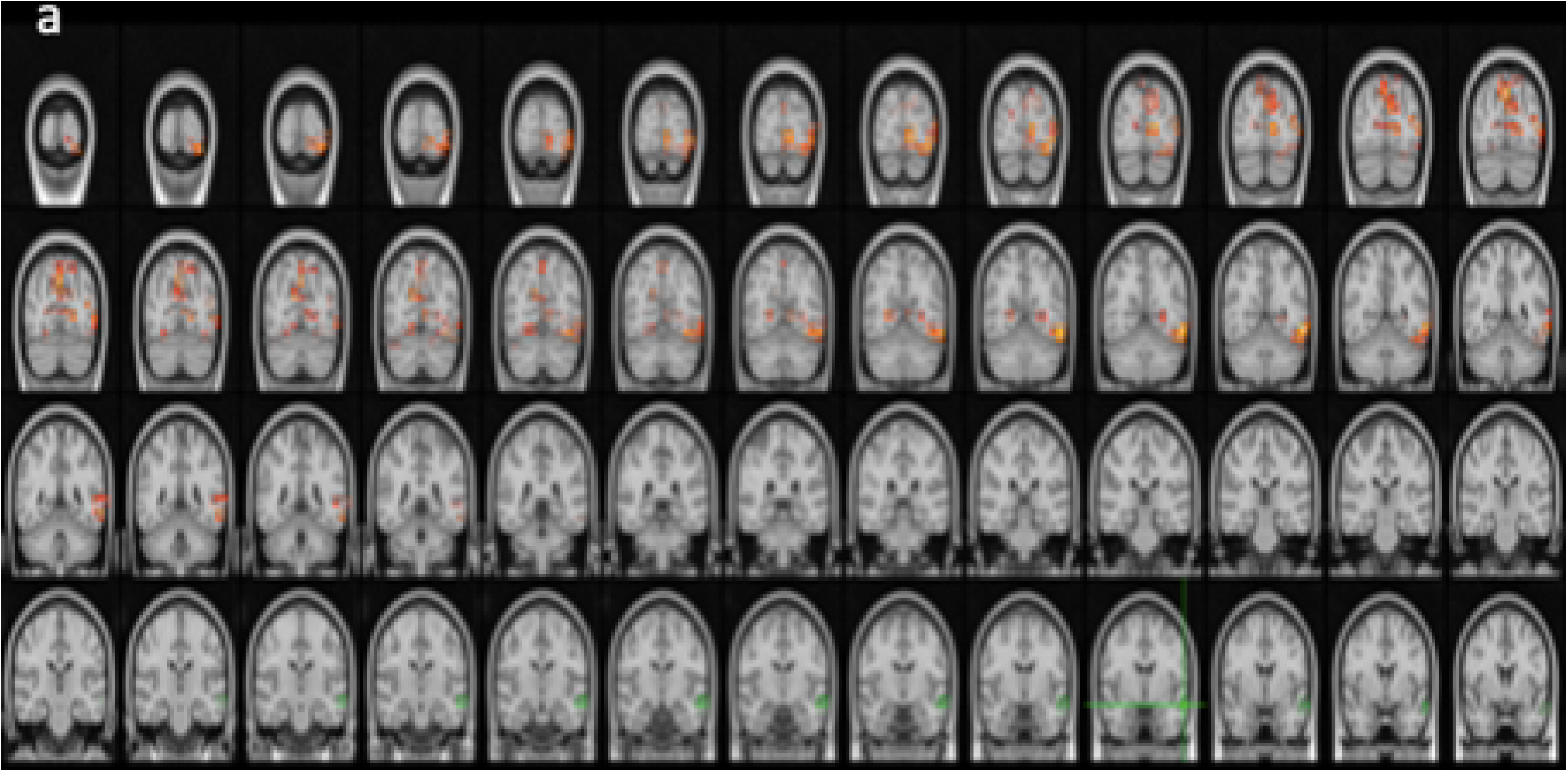
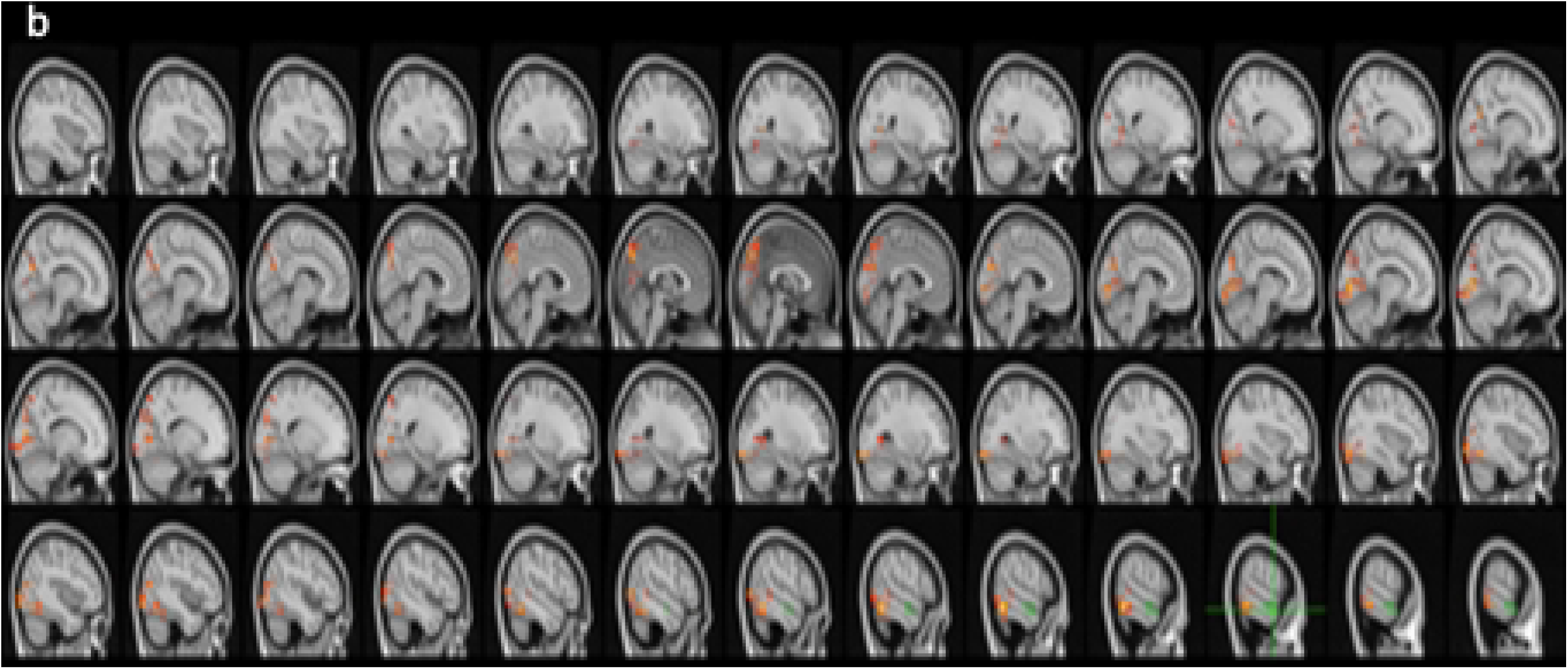
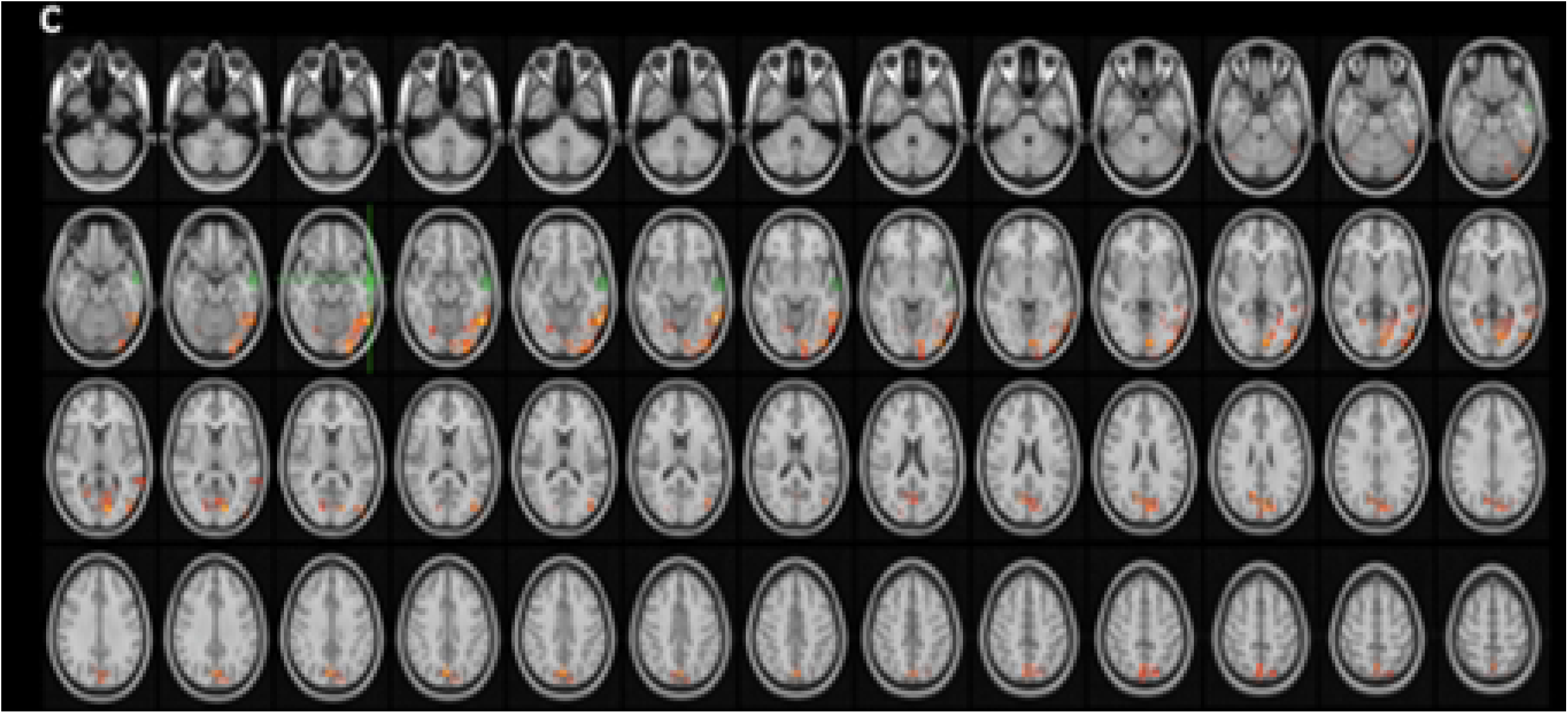
The areas with statistical-significantly increased functional connectivity between ROI in left medial temporal gyrus and bilateral occipital poles, bilateral occipital fusiform gyri, bilateral intra-calcarine cortices, right lingual gyrus, precuneus, medial temporal gyrus, when the time series of HS group and CBS(+) group are compared with one way t-test in dual regression analysis (p<0,05

**Fig 7.**
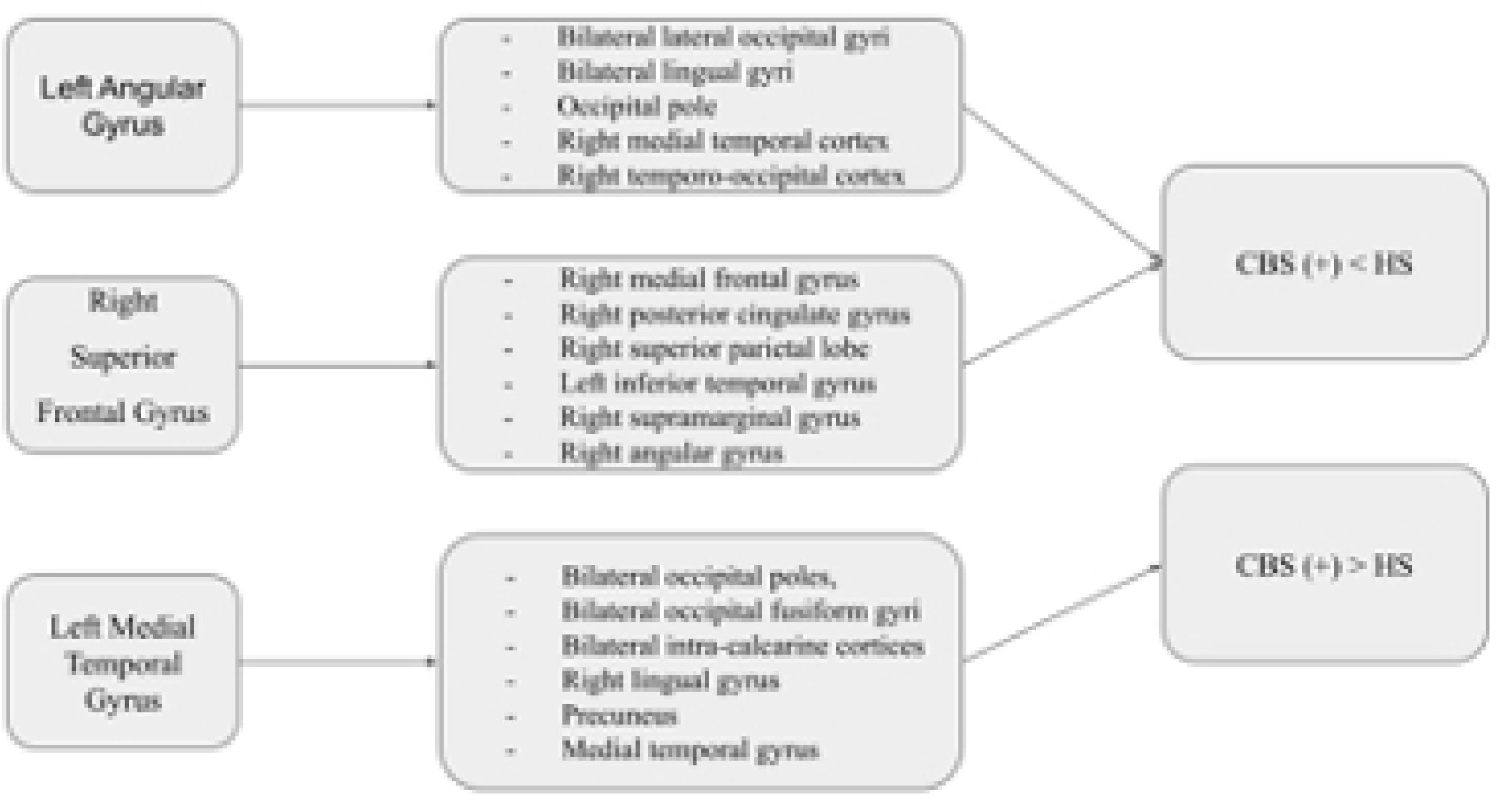
Summary of the Result

**Table 4.** The cluster coordinates with statistical-significantly increased functional connectivity between ROI in left medial temporal gyrus and bilateral occipital poles, bilateral occipital fusiform gyri, bilateral intra-calcarine cortices, right lingual gyrus, precuneus, medial temporal gyrus, when the time series of HS group and CBS(+) group are compared with one way t-test in dual regression analysis (p<0,05).

| Cluster Index | Voxels | 1-p-MAX | 1-p-MAX X (vox) | 1-p-MAX Y (vox) | 1-p-MAX Z (vox) | 1-p-COG X (vox) | 1-p-COG Y (vox) | 1-p-COG Z (vox) | COPE-MAX | COPE-MAX X (vox) | COPE-MAX Y (vox) | COPE-MAX Z (vox) | COPE-MEAN |
| --- | --- | --- | --- | --- | --- | --- | --- | --- | --- | --- | --- | --- | --- |
| Left medial temporal gyrus | 1847 | 0.99 | 74 | 36 | 31 | 63.3 | 26.3 | 33 | 0.99 | 74 | 36 | 31 | 964 |
| Precuneus | 638 | 981 | 43 | 26 | 55 | 45.5 | 25.6 | 53.5 | 981 | 43 | 26 | 55 | 962 |
| Right lingual gyrus | 67 | 964 | 37 | 30 | 31 | 34.9 | 28.5 | 30.2 | 964 | 37 | 30 | 31 | 955 |
| Left lateral occipital cortex | 56 | 961 | 52 | 26 | 62 | 53.5 | 26.1 | 61.3 | 961 | 52 | 26 | 62 | 955 |
| Left intra-calcarine cortex | 40 | 977 | 32 | 33 | 39 | 33.6 | 33 | 37.9 | 977 | 32 | 33 | 39 | 961 |
| Right intra-calcarine cortex | 40 | 961 | 37 | 26 | 40 | 36.8 | 23.2 | 41.7 | 961 | 37 | 26 | 40 | 953 |
| Left occipital fusiform gyrus | 15 | 956 | 62 | 29 | 33 | 62.5 | 29.9 | 34 | 956 | 62 | 29 | 33 | 953 |
| Left occipital fusiform gyrus | 13 | 956 | 57 | 30 | 29 | 57.5 | 29.1 | 28.8 | 956 | 57 | 30 | 29 | 953 |
| Left occipital fusiform gyrus | 12 | 0.96 | 56 | 23 | 25 | 55.6 | 22.9 | 25.2 | 0.96 | 56 | 23 | 25 | 954 |
| Right lateral occipital cortex | 6 | 953 | 24 | 29 | 24 | 24.5 | 29 | 23.5 | 953 | 24 | 29 | 24 | 952 |
| Right occipital fusiform gyrus | 3 | 955 | 53 | 29 | 25 | 52.7 | 29.3 | 25 | 955 | 53 | 29 | 25 | 953 |
| Left lateral occipital cortex | 1 | 953 | 61 | 27 | 51 | 61 | 27 | 51 | 953 | 61 | 27 | 51 | 953 |

## DISCUSSION

To the best of our knowledge, there have been few functional MRI studies evaluating the underlying pathophysiology of CBS hallucinations. Those studies that have been performed have included only a few patients, and due to their low sample sizes, their results are controversial. Most of the literature investigating CBS hallucinations is based on either metabolic (PET, SPECT) or electrophysiological (EEG) studies. However, this hurdle can be overcome with functional neuroimaging studies, including resting-state networks such as the default mode network, which is even sensitive to revealing overall brain activity during mind wandering and corresponding inner image generation. This may in turn strengthen the hypothesis of integrated visual hallucination and suggest a more generalized (plausible) mechanism underlying such hallucinations. Several pre-fMRI studies have revealed a decline in activity in primary and secondary visual cortices due to decreased visual perception from the outer world and increased activity in the heteromodal association areas from visual pathways, suggesting increased inner image generation. Interestingly, these studies also suggested decreased activity in frontal areas, even indicating a failure in the source monitoring system (3–5,7–9,28). For instance, a recent study investigating the structural and fMRI correlates of hallucinations in CBS patients and individuals with acquired blindness, and their corresponding healthy controls, revealed decreased gray matter volume in the medial occipital gyrus and cuneus in the CBS group and decreased gray matter volume in the medial occipital gyrus and lingual gyrus in the acquired blindness group compared to the healthy controls (29). The authors of that interesting work also suggested increased functional connectivity in the precuneus and secondary visual areas and decreased functional connectivity between the DMN and temporo-occipital fusiform gyrus in the CBS and blindness groups, respectively (29). These findings subsequently prompted researchers to investigate the underlying dynamic correlates of normal physiological states analogous to visual hallucinations, such as mind wandering in healthy conditions. In this context, when comparing fMRI parameters during visual hallucinations in CBS patients with complete visual loss and healthy controls during mind wandering (while watching visual images recreated based on the CBS patients’ visual hallucinations), Hahm et al. showed a special timely progressive pattern in the CBS group compared to the healthy controls. Based on that finding, the authors speculated that visual hallucinations are created similarly to Norhoff’s hypothesis for auditory hallucinations (30), suggesting that the visual pathway hierarchy autonomously creates visual hallucinations based on already existing inner created images (31).

Although our results concerning increased functional connectivity of the left medial temporal gyrus and primary visual areas with the precuneus and medial temporal lobe (two of the main hubs of the DMN) in the CBS(+) group compared to the healthy controls exhibit general agreement with Martial et al. and a previous metabolic study’s findings of increased functional connectivity in increased inner image creation, our findings are also in line with separate studies suggesting a deafferentationalist approach. For instance, we observed decreased functional connectivity in the CBS(+) group, not only between the angular gyrus (a heteromodal association hub that connects primary perception areas) and secondary visual cortices, but also with the right medial temporal and right temporo-occipital cortices (other heteromodal association areas). This may suggest decreased visual input and associated diminished functional connectivity between visual areas and heteromodal association areas in CBS(+) patients. It is also worth mentioning that our findings of decreased functional connectivity between the right superior frontal gyrus and heteromodal association areas (the main hubs for the DMN in CBS(+) patients) may provide further evidence for dysfunction in the source-monitoring system, since we observed decreased functional connectivity between specific front-rear structures of the DMN. It is important to emphasize here that instead of favoring one or other of these mechanisms, our findings confirm that these two components may be simultaneously active during hallucination generation, indicating a more holistic network model, as also previously suggested by Collerton et al. One of the main challenges we faced during our study was that the CBS(+) group participants were older, and female gender was more common in this group. A second limitation is that we found no statistically significant difference between the CBS(-) and healthy control groups, which may be due to the small sample size. However, this difficulty was irresoluble due to the lack of suitable CBS visual hallucination patients because of the rarity of the illness.

## CONCLUSION

Our findings suggest a combined mechanism in CBS related to increased internal created images caused by decreased visual external input causing visual hallucinations as well as impaired frontotemporal resource tracking system that together impair cognitive processing.

## ACKNOWLEDGEMENT

The authors of this work are grateful to Istanbul Medipol University, Research Institute for Health and Technologies (SABITA), Regenerative and Restorative Medicine Research Center (REMER), functional Imaging and Cognitive-Affective Neuroscience Lab (fINCAN).

## REFERENCES

1. Holroyd S, Rabins P V., Finkelstein D, Nicholson MC, Chase GA, Wisniewski SC. Visual hallucinations in patients with macular degeneration. Am J Psychiatry. 1992;149(12):1701–6.

2. Cogan DG. Visual hallucinations as release phenomena. Albrecht von Graefes Archiv fur klinische und experimentelle Ophthalmologie Albrecht von Graefe’s archive for clinical and experimental ophthalmology [Internet]. 1973 Jun [cited 2022 Jul 12];188(2):139–50. Available from: https://pubmed.ncbi.nlm.nih.gov/4543235/

3. Kazui H, Ishii R, Yoshida T, Ikezawa K, Takaya M, Tokunaga H, et al. Neuroimaging studies in patients with Charles Bonnet Syndrome. Psychogeriatrics [Internet]. 2009 [cited 2022 Jul 12];9(2):77–84. Available from: https://pubmed.ncbi.nlm.nih.gov/19604330/

4. Adachi N, Watanabe T, Matsuda H, Onuma T. Hyperperfusion in the lateral temporal cortex, the striatum and the thalamus during complex visual hallucinations: single photon emission computed tomography findings in patients with Charles Bonnet syndrome. Psychiatry Clin Neurosci [Internet]. 2000 [cited 2022 Jul 12];54(2):157–62. Available from: https://pubmed.ncbi.nlm.nih.gov/10803809/

5. Jang JW, Youn YC, Seok JW, Ha SY, Shin HW, Ahan SW, et al. Hypermetabolism in the left thalamus and right inferior temporal area on positron emission tomography-statistical parametric mapping (PET-SPM) in a patient with Charles Bonnet syndrome resolving after treatment with valproic acid. J Clin Neurosci [Internet]. 2011 Aug [cited 2022 Jul 12];18(8):1130–2. Available from: https://pubmed.ncbi.nlm.nih.gov/21700465/

6. Hanoglu L, Yildiz S, Polat B, Demirci S, Tavli A, Yilmaz N, et al. Therapeutic Effects of Rivastigmine and Alfa-Lipoic Acid Combination in the Charles Bonnet Syndrome: Electroencephalography Correlates. Curr Clin Pharmacol [Internet]. 2016 Oct 19 [cited 2022 Jul 12];11(4):270–3. Available from: https://pubmed.ncbi.nlm.nih.gov/27697039/

7. Ffytche DH. Visual hallucinatory syndromes: past, present, and future. Dialogues Clin Neurosci [Internet]. 2007 [cited 2022 Jul 12];9(2):173–89. Available from: https://pubmed.ncbi.nlm.nih.gov/17726916/

8. ffytche DH. The hodology of hallucinations. Cortex [Internet]. 2008 [cited 2022 Jul 12];44(8):1067–83. Available from: https://pubmed.ncbi.nlm.nih.gov/18586234/

9. Hanoglu L, Yildiz S, Cakir T, Hanoglu T, Yulug B. FDG-PET Scanning Shows Distributed Changes in Cortical Activity Associated with Visual Hallucinations in Eye Disease. Endocr Metab Immune Disord Drug Targets [Internet]. 2019 Aug 30 [cited 2022 Jul 12];19(1):84–9. Available from: https://pubmed.ncbi.nlm.nih.gov/30160221/

10. Collerton D, Perry E. Dreaming and hallucinations - continuity or discontinuity? Perspectives from dementia with Lewy bodies. Conscious Cogn [Internet]. 2011 Dec [cited 2022 Jul 12];20(4):1016–20. Available from: https://pubmed.ncbi.nlm.nih.gov/21531149/

11. Raichle ME, MacLeod AM, Snyder AZ, Powers WJ, Gusnard DA, Shulman GL. A default mode of brain function. Proc Natl Acad Sci U S A [Internet]. 2001 Jan 16 [cited 2022 Jul 12];98(2):676–82. Available from: www.pnas.org

12. Buckner RL, Andrews-Hanna JR, Schacter DL. The brain’s default network: anatomy, function, and relevance to disease. Ann N Y Acad Sci [Internet]. 2008 Mar [cited 2022 Jul 12];1124:1–38. Available from: https://pubmed.ncbi.nlm.nih.gov/18400922/

13. Qin P, Northoff G. How is our self related to midline regions and the default-mode network? Neuroimage [Internet]. 2011 Aug 1 [cited 2022 Jul 12];57(3):1221–33. Available from: https://pubmed.ncbi.nlm.nih.gov/21609772/

14. Delli Pizzi S, Franciotti R, Tartaro A, Caulo M, Thomas A, Onofrj M, et al. Structural alteration of the dorsal visual network in DLB patients with visual hallucinations: a cortical thickness MRI study. PLoS One [Internet]. 2014 Jan 23 [cited 2022 Jul 12];9(1). Available from: https://pubmed.ncbi.nlm.nih.gov/24466177/

15. Shine JM, O’Callaghan C, Halliday GM, Lewis SJG. Tricks of the mind: Visual hallucinations as disorders of attention. Prog Neurobiol [Internet]. 2014 [cited 2022 Jul 12];116:58–65. Available from: https://pubmed.ncbi.nlm.nih.gov/24525149/

16. Onofrj M, Taylor JP, Monaco D, Franciotti R, Anzellotti F, Bonanni L, et al. Visual hallucinations in PD and Lewy body dementias: old and new hypotheses. Behavioural neurology [Internet]. 2013 [cited 2022 Jul 12];27(4):479–93. Available from: https://pubmed.ncbi.nlm.nih.gov/23242366/

17. Muller AJ, Shine JM, Halliday GM, Lewis SJG. Visual hallucinations in Parkinson’s disease: theoretical models. Mov Disord [Internet]. 2014 Nov 1 [cited 2022 Jul 12];29(13):1591–8. Available from: https://pubmed.ncbi.nlm.nih.gov/25154807/

18. Franciotti R, Delli Pizzi S, Perfetti B, Tartaro A, Bonanni L, Thomas A, et al. Default mode network links to visual hallucinations: A comparison between Parkinson’s disease and multiple system atrophy. Mov Disord [Internet]. 2015 Aug 1 [cited 2022 Jul 12];30(9):1237–47. Available from: https://pubmed.ncbi.nlm.nih.gov/26094856/

19. Yao N, Shek-Kwan Chang R, Cheung C, Pang S, Lau KK, Suckling J, et al. The default mode network is disrupted in Parkinson’s disease with visual hallucinations. Hum Brain Mapp [Internet]. 2014 Nov 1 [cited 2022 Jul 12];35(11):5658–66. Available from: https://pubmed.ncbi.nlm.nih.gov/24985056/

20. Bejr-kasem H, Pagonabarraga J, Martínez-Horta S, Sampedro F, Marín-Lahoz J, Horta-Barba A, et al. Disruption of the default mode network and its intrinsic functional connectivity underlies minor hallucinations in Parkinson’s disease. Mov Disord [Internet]. 2019 Jan 1 [cited 2022 Jul 12];34(1):78–86. Available from: https://pubmed.ncbi.nlm.nih.gov/30536829/

21. Yildiz S, Yulug B, Kocabora MS, Hanoglu L. Power spectral density and coherence analysis of eye disease with and without visual hallucination. Neurosci Lett [Internet]. 2021 Jan 1 [cited 2022 Jul 12];740. Available from: https://pubmed.ncbi.nlm.nih.gov/33127444/

22. Güngen C, Ertan T, Eker E, Yaşar R, Engin F. [Reliability and validity of the standardized Mini Mental State Examination in the diagnosis of mild dementia in Turkish population] - PubMed. Turk Psikiyatri Derg [Internet]. 2002 [cited 2022 Jul 12];13(4):273–81. Available from: https://pubmed.ncbi.nlm.nih.gov/12794644/

23. Papapetropoulos S, Katzen H, Schrag A, Singer C, Scanlon BK, Nation D, et al. A questionnaire-based (UM-PDHQ) study of hallucinations in Parkinson’s disease. BMC Neurol [Internet]. 2008 Jun 20 [cited 2022 Jul 12];8. Available from: https://pubmed.ncbi.nlm.nih.gov/18570642/

24. Yesavage JA, Brink TL, Rose TL, Lum O, Huang V, Adey M, et al. Development and validation of a geriatric depression screening scale: a preliminary report. J Psychiatr Res [Internet]. 1982 [cited 2022 Jul 12];17(1):37–49. Available from: https://pubmed.ncbi.nlm.nih.gov/7183759/

25. Brown CA, Schmitt FA, Smith CD, Gold BT. Distinct patterns of default mode and executive control network circuitry contribute to present and future executive function in older adults. Neuroimage [Internet]. 2019 Jul 15 [cited 2022 Jul 12];195:320–32. Available from: https://pubmed.ncbi.nlm.nih.gov/30953834/

26. JASP Team. JASP (Version 0.14.1)[Computer software]. 2020.

27. George D, Paul Mallery with. SPSS for Windows Step by Step A Simple Guide and Reference Fourth Edition (11.0 update) Answers to Selected Exercises.

28. Graham G, Dean J, Mosimann UP, Colbourn C, Dudley R, Clarke M, et al. Specific attentional impairments and complex visual hallucinations in eye disease. Int J Geriatr Psychiatry [Internet]. 2011 Mar [cited 2022 Jul 12];26(3):263–7. Available from: https://pubmed.ncbi.nlm.nih.gov/20684031/

29. Martial C, Larroque SK, Cavaliere C, Wannez S, Annen J, Kupers R, et al. Resting-state functional connectivity and cortical thickness characterization of a patient with Charles Bonnet syndrome. PLoS One [Internet]. 2019 Jul 1 [cited 2022 Jul 12];14(7). Available from: https://pubmed.ncbi.nlm.nih.gov/31318888/

30. Northoff G. Are Auditory Hallucinations Related to the Brain’s Resting State Activity? A “Neurophenomenal Resting State Hypothesis.” Clin Psychopharmacol Neurosci [Internet]. 2014 Dec 1 [cited 2022 Jul 12];12(3):189–95. Available from: https://pubmed.ncbi.nlm.nih.gov/25598821/

31. Hahamy A, Wilf M, Rosin B, Behrmann M, Malach R. How do the blind “see”? The role of spontaneous brain activity in self-generated perception. Brain [Internet]. 2021 Jan 1 [cited 2022 Jul 12];144(1):340–53. Available from: https://pubmed.ncbi.nlm.nih.gov/33367630/

